# Interpreting Protein Language Models: high attention sites predict functional regions

**DOI:** 10.64898/2026.07.29.741641

**Authors:** Sophia J. Pribus, Russ B. Altman, Gowri Nayar

## Abstract

Computational proteomics has revolutionized biomedical research, guiding targeted experimental exploration to accelerate protein-based mechanistic discovery. Protein Language Models (PLMs) enable scalable, resource-efficient study; through large-scale training on only primary protein sequences, PLMs generate vector representations of protein structure that have been shown to capture biochemical, evolutionary, and structural properties. A core component of PLMs is the attention mechanism, which specifically captures long-range interactions across a protein sequence in attention matrices. Using the Evolutionary Scale Modelling 2 (ESM-2) PLM, we previously developed a novel method to identify “High Attention” (HA) sites. HA sites are specific residues that ESM-2 assigns the most attention to early during encoding. Here, we further characterize these HA sites across structural and functional metrics. Using unsupervised clustering, we find HA sites can be categorized as “structural core”, “structural pathogenic”, “core pathogenic”, or “low-confidence”. We further use AlphaMissense pathogenicity predictions and the pan-cancer analysis of whole genomes (PCAWG)-labeled pathogenic variant positions to show that HA sites predict protein regions with high pathogenic risk. Finally, we explore the utility of HA sites for suggesting candidate binding sites, identifying multiple cancer protein examples where HA sites identified regions with previously undiscovered high interaction likelihood and thus potential therapeutic utility. Our work demonstrates the biological interpretability of PLM representations and offers a valuable method to prioritize functionally relevant protein residues for targeted biomedical research.

## 1. Introduction

Linking protein structure to function is crucial for biomedical tasks, including disease characterization and drug development.^1^ However, while the human proteome is composed of over 20,000 proteins, we have experimental characterization of just 24% of residues across these proteins.^2^ Practical experimental challenges, including innate protein structure instability and the expensive, time-consuming nature of functional and structural determination procedures, have hindered systematic proteomics analysis.^3^ Computational proteomics tools, including structure prediction models (e.g. AlphaFold^4^) and protein language models (e.g. Evolutionary Scale Modelling [ESM]^5^), have revolutionized our ability to study proteins.

Central to our research is the understanding that protein structure drives function.^6^ Protein activity alterations drive pathogenesis across the spectrum of disease, and understanding how, when, and why these alterations occur is an essential step in disease characterization.^1^ For example, the precise 3D fold of a protein defines regions where the protein may engage in catalytic reactions with other proteins, or “active sites”. Similarly, the 3D fold defines areas where the protein may form strong links with other molecules, including drugs, or “binding sites”. Certain alterations to the 3D structure, through mutational or inhibitory processes, may cause impaired protein function and thus disease; thus, it is valuable to study the “pathogenicity” of certain protein areas (i.e., how likely they are to cause dysfunction and disease, if modified).

Computational tools enable rapid, scalable exploration of protein structures. Protein Language Models (PLMs), specifically, apply the transformer-based mechanisms originally designed for natural language processing to protein sequence.^7^ In pre-training, PLMs take protein primary sequences as input and are trained to predict masked residues. This process produces internal representations, called embeddings, which capture biochemical characteristics of individual amino acids and residue interactions that dictate local and global structure and function.^8^ A key component of PLM encoding is their attention mechanisms, which capture these residue interactions. PLMs use multiple layers of attention; at each layer, each residue is given a weighted attention score based on their interactions with other residues in the sequence.^7,9^ Thus, the resulting attention matrices indicate the relative importance of each residue at each layer of the encoder, highlighting those residues which emerge early as key sequence positions.

Prior methods^9^ have shown that residues with high attention values have significant links to protein structure and function. High attention residues were found to localize near protein active sites, demonstrating that PLM attention matrices have functional interpretations. Additionally, high attention residues were found to be conserved across protein families, showing that PLM attention matrices encode structural patterns across proteins.^9^ While these are strong indicators of the biological interpretability of PLM attention matrices, there is still a need to fully explore how attention patterns correlate with other structural and functional features, like bindability and pathogenicity. There is also a question of whether additional residues emerging at deeper attention matrix layers than the original high attention sites, but still early in encoding, have similar structural and functional links. We thus introduce the idea of primary, secondary and tertiary HA sites, with primary HA sites being those emerging at the earliest layer, followed by secondary and tertiary HA sites in subsequent layers.

In this work, we extend previous methods^9^ to identify primary, secondary, and tertiary high attention sites, discovered sequentially from the ESM-2 attention matrices.^10^ We show that high attention sites correlate with other key structural and functional metrics, including known binding sites and protein domains and residues with high mutational pathogenicity. We thus expand the biological interpretability of PLMs and demonstrate the utility of high attention sites for efficient, targeted biomedical proteomics experiments.

## 2. Method

We identify high attention (HA) sites across the human proteome and characterize these residues across structural and functional metrics. Building on our initial work, we use unsupervised learning to categorize HA site types and demonstrate their utility to prioritize functionally relevant positions for biomedical research. An overview of our complete method is visualized in **Figure 1**.

**Fig 1.**
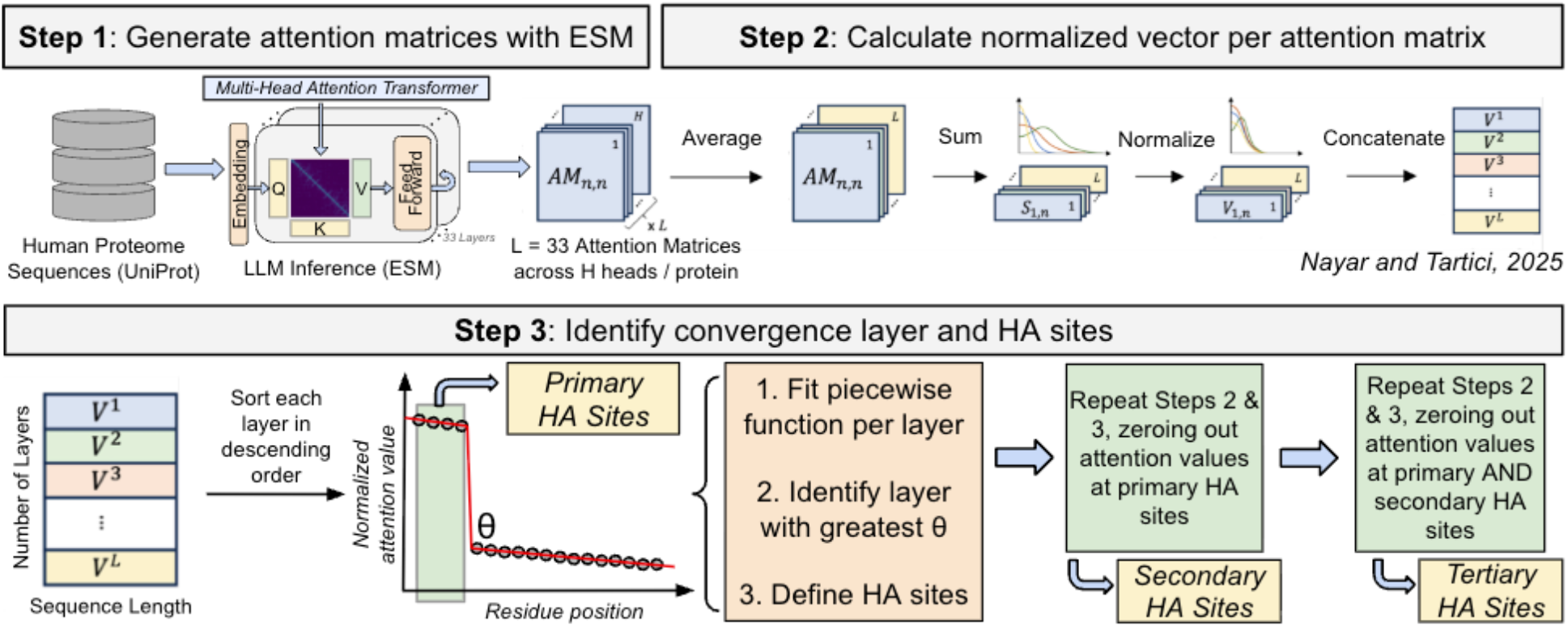
Overview of method to define primary, secondary, and tertiary high attention sites. We show a high-level visualization of the method used to identify all high attention (HA) sites, the subset of residues which emerge at the earliest ESM-2 layer as having high attention across pooled attention heads. In Step 1, we use the ESM model to analyze all human proteins in the UniProt database, running the protein language model on primary sequences and extracting attention matrices for each protein. In Steps 2-3, we apply our method (described in **methods 2.1**) and obtain “primary” HA sites, the first emerging high attention sites. We then obtain secondary and tertiary HA sites by starting with the attention vectors derived as usual at the end of step 2 but replacing the attention values at the previously identified primary HA site indices with 0. We then apply the same step 3 method to these updated attention vectors to derive “secondary” HA sites. “Tertiary” HA sites are identified similarly, replacing both primary and secondary HA site indices with 0 in the attention vectors at the end of step 2.

### 2.1. Defining high attention (HA) sites from PLM attention matrices

ESM-2 is given primary protein sequences as input and trained to predict masked residues. To do so, it uses a transformer mechanism based on self-attention, learning the interactions between pairwise residues across the span of the protein sequence. ESM-2 generates attention matrices (n-by-n, where n = number of residues in the protein sequence) across multiple heads (in our study, 14 attention heads). It trains through numerous layers (in our study, 33 layers), starting from a randomized attention input and progressively fine-tuning pairwise interactions.

Previously, for each protein we summarized the attention at each residue across layers by averaging attention across all attention heads, summing the attention at each residue across all pairwise interactions, and normalizing the attention values across the length of the protein sequence (**Figure 1**, steps 1-2).^9^ We then generated normalized attention heatmaps by displaying these attention vectors organized by layer, from 1 to 33 (**Figure 1**, step 2; **Figure 2**). We noted a distinct layer at which specific residues acquire significantly concentrated pooled attention relative to other residues (**Figure 2**, pink box).^9^ These were defined conceptually as “high attention” residues, i.e. the subset of residues which emerge at the earliest layer as having high attention across pooled attention heads. In this study, we re-define these as “primary” HA sites, to indicate their status as the first emerging high attention sites.

**Fig 2.**
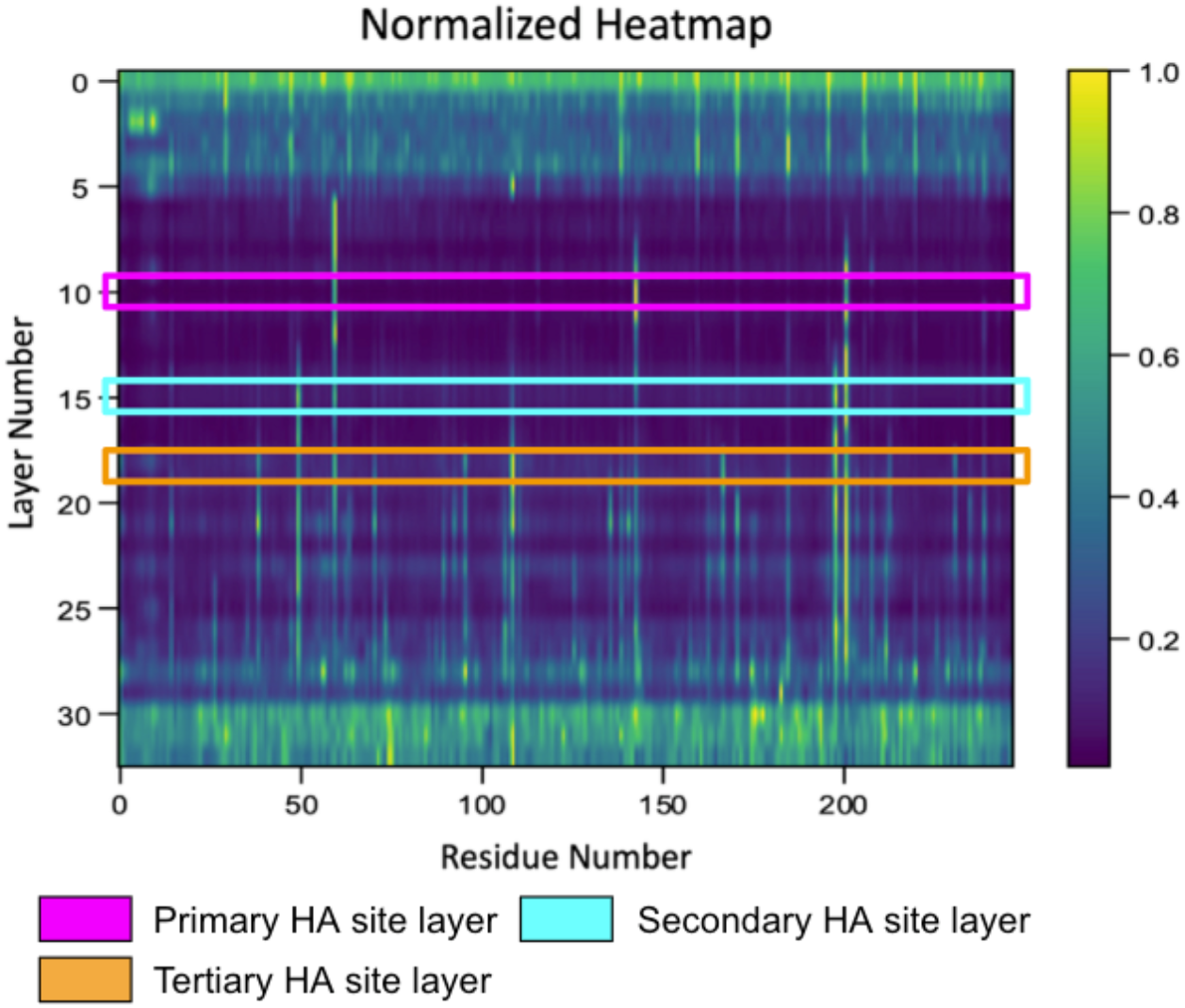
Defining primary, secondary, and tertiary HA sites from normalized attention heatmap. For protein P07477, normalized attention heatmap is generated by displaying attention vectors organized by layer, from 0 to 32 (33 total layers), as described in **Figure 1**. Boxes indicate the layers where successive high attention residues are identified: pink for primary HA site layer, cyan for secondary HA site layer, and orange for tertiary HA site layer.

### 2.2. Identifying primary, secondary, and tertiary HA sites

After calculating primary HA sites as described in our previous publication^9^ and above, we note that in the layers following the layer at which primary HA sites are identified, several additional residues emerge with high attention; we define these as “secondary” HA sites (**Figure 2**, cyan box). To calculate secondary HA sites, we start with the normalized and pooled attention value vectors *v*_*l*_ for each layer *l*. We replace the attention values at the previously identified primary HA site indices with 0, to ensure that the same residues are not identified. We then use the same original high-attention identification method with the updated attention values but only consider attention layers at least three layers deeper than the primary convergence layer (i.e., the convergence layer at which primary HA sites were identified) (**Figure 1**). We add this layer buffer to avoid selecting secondary HA sites highly proximal to primary HA sites; searching only in layers three layers beyond the primary convergence layer returns HA sites at regions distant from primary sites. In addition, residues close to the original high attention sites are likely highly attended to as an artifact of the cross-attention model, representing the localized importance surrounding the original primary HA site. The buffer of n = 3 layers was chosen based on visual assessment of concatenated attention vectors, noting that new attention “hotspots” appeared to surface around three layers after the initially identified convergence layer (case example shown in **Figure 2**).

As done in our previous publication to calculate primary HA sites, we then sort each subsequent *v*_*l*_ in descending order, fitting linear segments to the curve, calculating the angle θ between the linear segments, and selecting the layer with θ closest to 90 degrees as the convergence layer.^9^ Residues to the left side of the breakpoint are selected as the secondary HA sites. We similarly calculate “tertiary” HA sites (**Figure 2**, orange box), replacing attention values at the previously identified primary and secondary HA site indices with 0 and continuing with the method for attention layers at least five layers deeper than the secondary convergence layer (**Figure 1**). This layer buffer of n = 5 layers was similarly chosen based on visual assessment of concatenated attention vectors. We chose to stop at tertiary HA sites as the remaining layers tended to display high attention across more than half of the residues, reducing the interpretability of specific HA site qualities. For clarity and brevity, we include the stratified results per level of HA site in the supplement only.

### 2.3. Calculating structural and functional metrics at HA sites

#### Active sites

For proteins with annotated active sites, we obtain all active site residues from UniProt.^11^ Using AlphaFold2-generated three-dimensional protein structures, we calculate the spatial distance in Angstroms from each HA site residue to the nearest active site residue, defined as the Minimum Spatial Distance (MSD).

#### Binding sites

For proteins with annotated binding sites, we obtain all binding site residues from UniProt. Using AlphaFold2-generated three-dimensional protein structures, we calculate MSD from each HA site to the nearest binding site. For an extended set of binding sites, we also obtained binding sites from BioLIP^12^ and similarly calculated the MSD from each HA site to the nearest binding site.

#### Domain sites

For proteins with annotated domains, specific combinations of secondary structures organized into three-dimensional folds^13^, we obtain all domain residues from UniProt. Using AlphaFold2-generated three-dimensional protein structures, we calculate MSD from each HA site to the nearest protein domain.

#### Centrality

We calculate the radial position of each HA site relative to the protein centroid. We find the protein centroid by calculating the three-dimensional average of all atom coordinates in the AlphaFold2-generated three-dimensional structure. We then calculate the spatial distance from each HA site’s central carbon atom to the protein centroid. We scale this distance by dividing by the maximum distance from the protein centroid to any atom in the AlphaFold2-generated structure, reporting a scaled (0-1) HA position within the protein.

#### AlphaFold score

We report AlphaFold2’s structural prediction certainty score, the predicted local distance difference test (pLDDT), at each HA site.^4^

#### Pathogenicity metrics

To assess pathogenicity at HA sites, we use AlphaMissense^14^ pathogenicity scores.^15^ We calculate a pathogenicity score at each HA site by averaging the AlphaMissense pathogenicity scores across all mutations at that residue. In addition, for each protein, we calculate the top 10 pathogenic sites by averaging the pathogenicity scores for each mutation at each residue and selecting the 10 sites with the highest average. We then calculate the MSD from each HA site to the nearest of these 10 sites.

#### Amino acid and secondary structure type

Using DSSP^16^, we extract the amino acid type of each HA site and record any secondary structure type the HA site is found within.

#### Randomization comparisons

We generate two sets of random sites for all statistical comparisons with true HA sites. The random site sets are calculated per protein and are matched in number to the original count of HA sites for each protein. We calculate distinct random sites for primary, secondary, and tertiary HA sites. The first set is <u>pure random sites</u>, selected without replacement from all non-HA site residues in the protein. The second set is <u>residue-matched (RM) random sites</u>, selected from all non-HA site residues in the protein that are the same amino acid type as the HA sites. For example, if the HA sites for a specific protein are glycine and valine residues, then the first RM random site would be selected from all non-HA site glycine residues and the second from all non-HA site valine residues. We compare all metric distributions to these pure random and residue-matched (RM) random residue sets, using two-sided Mann-Whitney tests to assess pairwise statistical significance. For amino acid frequency and secondary structure frequency, we use Chi-squared analysis.

### 2.4. Categorizing HA site types with unsupervised clustering

We cluster HA sites using the K-means unsupervised clustering method. We generate an elbow plot of the number of clusters against clustering inertia to select K for each clustering. We clustered primary, secondary, and tertiary HA sites both independently and grouped together. We clustered based on a reduced set of metrics, including only AlphaFold score, AlphaMissense average score, and normalized distance from protein centroid. We calculate the Pearson correlation between these metrics and find only modest correlations between each pair (**Supplementary fig. 1a**), indicating they are sufficiently unique to include as distinct features in clustering. We use an elbow plot with K-means clustering to determine the optimal number of clusters, finding it to be K = 4 (**Supplementary fig. 1b**). From this clustering, we identified four categories of HA sites:

1. Structural core HA sites - sites with low AlphaMissense average pathogenicity scores, proximal to the protein centroid.
2. Structural pathogenic HA sites - sites with high AlphaMissense average pathogenicity scores, at the mid-region between protein centroid and surface.
3. Core pathogenic HA sites - sites with high AlphaMissense average pathogenicity scores, proximal to the protein centroid.
4. Low-confidence HA sites - sites with low AlphaFold prediction confidence scores.

To assess the robustness of these categories, we similarly perform unsupervised clustering with the subset of human proteins with experimentally-determined active sites, binding sites, and domains, for which we calculate a larger set of metrics, including minimum spatial distance (MSD) to active sites, MSD to binding sites, MSD to protein domains, MSD to AlphaMissense top 10 pathogenic sites, AlphaFold prediction confidence score, AlphaMissense average pathogenicity score, protein sequence length, and normalized distance from protein centroid. We calculate the pairwise correlation between these metrics and similarly found only modest correlation between pairs (**Supplementary fig. 1c**). We use an elbow plot with K-means clustering and again found the optimal number of clusters to be K = 4 (**Supplementary fig. 1d**). In all clustering methods, any HA sites with incomplete metrics were excluded from clustering.

### 2.5. Evaluating relationship of HA sites with documented pathogenic variants

In addition to calculating the AlphaMissense average pathogenicity scores at each HA site, we calculate the distance from each HA site to known pathogenic variant positions. We obtain variant positions from the pan-cancer analysis of whole genomes (PCAWG) dataset^17^, sub-setting to missense, nonsense, and silent mutation types. We then calculate the minimum and average MSD from each HA site to each mutation.

### 2.6. Evaluating use of HA sites for binding site discovery

We select a sample of classically “undruggable” proteins, including MYC and BCL-2 and identify their HA sites. We define a region of 4-8 Angstroms around these HA sites. We use fPocket^18^ to assess the druggability of these regions, reporting the druggability probability.

## 3. Results

### 3.1. HA sites are proximal to known binding sites and have high predicted pathogenicity scores

Across grouped primary, secondary, and tertiary HA sites, we observe that HA sites are significantly closer to active sites, binding sites, protein domains, and protein centroids than random residues (**Figure 3, Supplementary figs. 2-4**), demonstrating they track with key structural areas of proteins, including areas with functional significance (i.e., active and binding sites). We observe high AlphaFold^4^ predication confidence scores with a notable left skew of low scores, suggesting most HA sites are found at structurally common sites, with a subset found in unique 3D arrangements poorly represented by AlphaFold’s training data. We observe significantly increased AlphaMissense average pathogenicity scores compared to random residues (**Figure 3, Supplementary figs. 2-4**), indicating HA sites support healthy protein function and lead to pathogenic states when altered.

**Fig 3.**
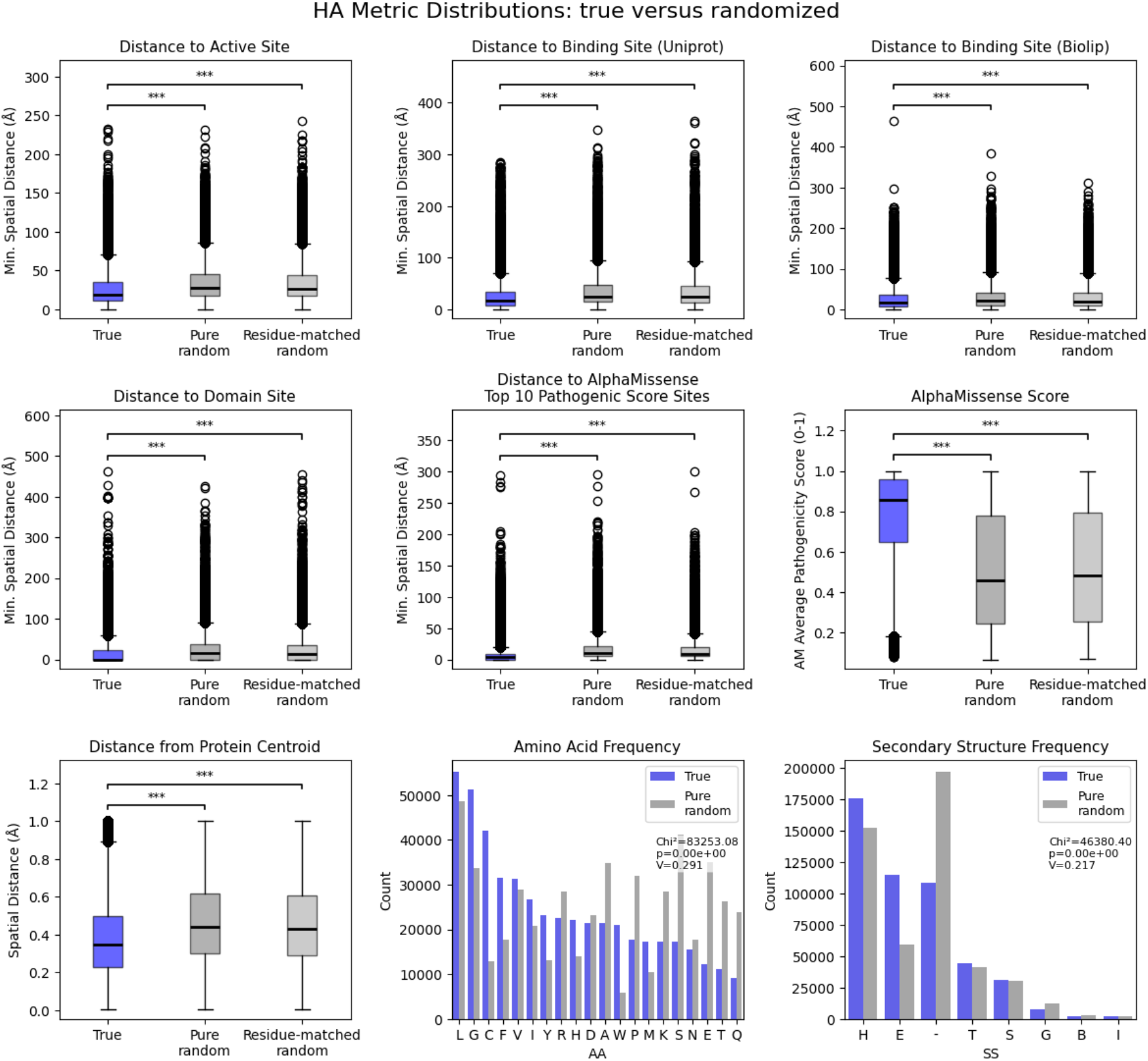
High attention site structural and functional metric distributions. To evaluate the biological relevance of the HA sites, we plot structural and functional metrics of the pooled primary, secondary, and tertiary HA sites (**methods 2.3**). Structural metrics include spatial orientation of the HA sites, described by the minimum spatial distance from HA sites to known active sites, binding sites, and domain sites (from UniProt and BioLIP documentations), as well as the calculated protein centroid. Structural metrics also include HA site primary and secondary structure, shown as frequency plots of the amino acid type of HA sites and the secondary structures HA sites are part of. Functional metrics focus on pathogenicity of HA sites as predicted by AlphaMissense; the minimum distance from HA sites to the top 10 pathogenic sites in the protein as predicted by AlphaMissense, as well as the AlphaMissense score at the HA site, are shown. In blue we plot the metrics calculated at true HA sites. In dark grey, we plot the metrics calculated at randomized positions in the protein. In light grey, we plot the metrics calculated at residue-matched random positions (i.e., positions randomly selected from all residues of the same amino acid type as a given HA site). Two-sided Mann-Whitney tests with Bonferroni correction and Chi-squared analysis used to assess statistical significance. *** indicates p-value < 0.001. V = Cramer’s V. Amino acid single letter codes match standard definitions. Secondary structure single letter codes are defined as follows: E = extended strand, participates in β ladder; H = α-helix; - = coil / irregular (no defined secondary structure); T = hydrogen-bonded turn, S = bend, P = κ-helix (poly-proline II helix), G = 3_10_-helix; I = π-helix; B = residue in isolated β-bridge. Across structural and functional metrics, we highlight the proximity of true HA sites to active and binding sites, as well as their significantly enriched pathogenicity (as calculated by the AlphaMissense score).

### 3.2. Unsupervised clustering defines four distinct categories of HA sites

Considering the distribution tails across structural and functional metrics, we next explore whether specific categories of HA sites can be defined according to patterns in where they land along different metric distributions. Based on the clustering result presented in **figure 4**, we define four categories of HA sites: low-confidence, structural core, structural pathogenic, core pathogenic (see **Methods 2.4** for definitions).

**Fig 4.**
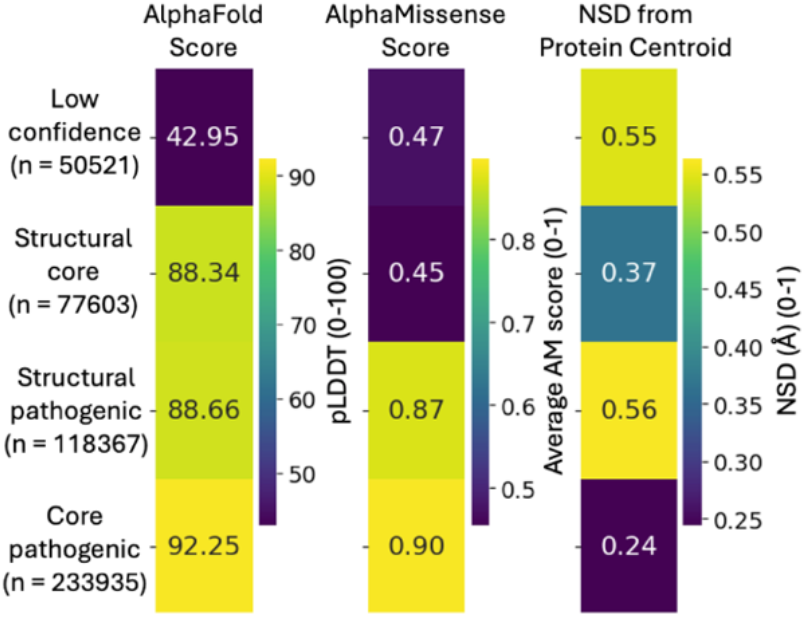
Unsupervised clustering reveals four categories of HA sites. Clustering performed on metrics (AlphaFold score, AlphaMissense score, NSD from protein centroid) calculated per HA site (pooled primary, secondary, and tertiary). K-means clustering was used, returning an optimal K = 4 (**methods 3.2**). AlphaFold Score = predicted local distance difference test (pLDDT) score; AlphaMissense Score = pathogenicity score; NSD from Protein Centroid = normalized spatial distance in Angstroms from protein centroid. Values in box represent average metric value for cluster. Left column counts represent number of HA sites in each cluster. This clustering reveals four distinct categories of HA sites, defined by their spatial characterization within the protein and functional value, as defined by the AlphaMissense score.

### 3.3. Structural pathogenic and core pathogenic HA sites are enriched near documented pathogenic variants

We find that structural pathogenic and core pathogenic HA sites have high predicted pathogenicity scores, as expected by their definitions (**Figure 5**). In a real-world, clinical context, we further observe that HA sites are significantly closer to the pan-cancer analysis of whole genomes (PCAWG) database variants compared to random residues (**Figure 6a**), a trend that holds when splitting by variant type (missense, nonsense, or silent) and across independent HA site types (primary, secondary, and tertiary) (**Supplementary fig. 9a**). Thus, in both computational predictions and clinical cohorts, HA sites are enriched at and near positions where mutations are pathogenic, demonstrating their clinical relevance.

**Fig 5.**
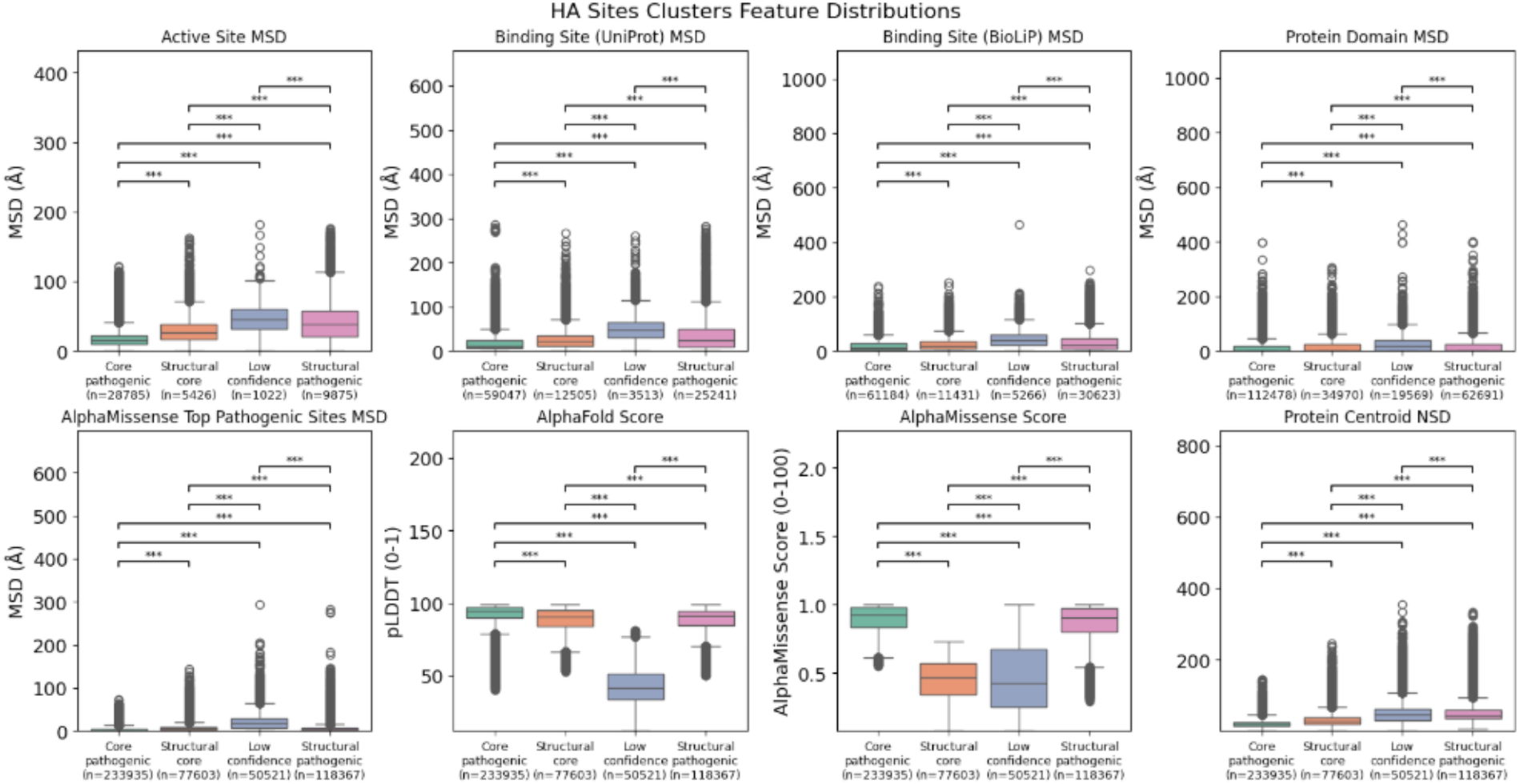
HA site structural and functional metrics by cluster. To evaluate the broader biological distinctions of the four categories of HA sites, we plot the calculated structural and functional metrics by cluster (**methods 3.3, 3.4**). Two-sided Mann-Whitney tests with Bonferroni correction used to assess statistical significance. *** indicates p-value < 0.001. We note significant variation across the groups with distinct trends, indicating that HA site categories may be used to suggest HA sites with specific structural and functional features (e.g. binding site proximity, pathogenicity, etc.).

**Fig 6.**
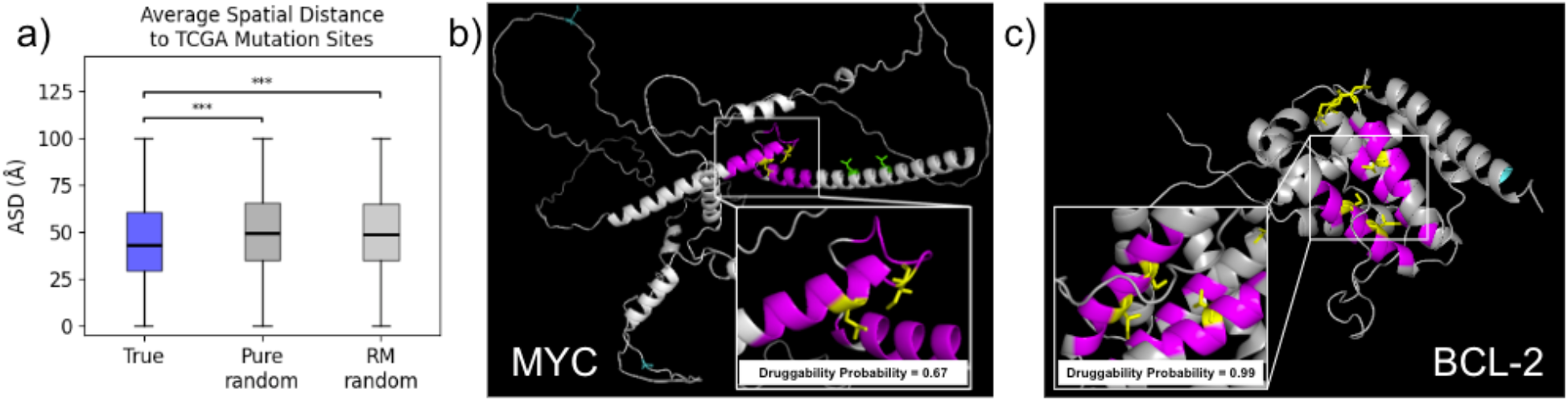
HA sites are enriched near pathogenic variants and define novel binding sites. **a)** Boxplot shows the average spatial distance (ASD) in Angstroms between each HA site residue and the nearest pathogenic variant (missense, nonsense, or silent) documented in the PCAWG cohort. Calculated distribution is plotted in blue and compared to both sets of randomized sites (**methods 2.3, 2.5**). In dark grey, we plot the metrics calculated at randomized positions in the protein. In light grey, we plot the metrics calculated at residue-matched random positions. Two-sided Mann-Whitney tests with Bonferroni correction used to assess statistical significance. *** indicates p-value < 0.001. **b-c)** PyMOL visualization of AlphaFold-generated structures of the MYC protein (**b**) and BCL-2 protein (**c**) with HA sites highlighted (**methods 3.6**). Yellow = core pathogenic HA sites, green = structural pathogenic HA sites, cyan = low-confidence HA sites. Pink = pocket of 8Å around core pathogenic HA sites. fPocket-predicted pocket druggability probability is shown for single pocket defined by core pathogenic HA sites (1 pocket / 3 total identified for BCL-2, see others in **supplementary figure 9**); fPocket defines threshold of 0.5 for probable bindability of the pocket, probability near 1 indicates high likelihood of molecular binding.

### 3.4. Using core HA sites to predict candidate binding sites

We find structural core HA sites and core pathogenic HA sites (defined in **Methods 2.4**) are more proximal to active sites and binding sites than are the other two categories of HA sites (**Figure 5**). We explore whether these HA sites can be used to define candidate binding positions for proteins without known binding sites. In both MYC and BCL-2, classically undruggable proteins^19^, we found new binding regions with druggability probabilities above 0.5 (as predicted by fPocket^18^; see **Methods 2.6**). For MYC, we found one binding region with druggability probability equal to 0.67

(**Figure 6b**); this region has previously been shown to bind numerous compounds in vitro with inhibitory effects on cell cycle progression.^20^ For BCL-2, we found three binding regions with druggability probabilities of 0.99 (**Figure 6c**), 0.91 (**Supplementary fig. 9b**), and 0.65 (**Supplementary fig. 9c**). These regions are also supported as potential binding sites by prior literature, including the traditionally targeted BH3 domain^21,22^, the previously suggested BH4 domain target^22^, and the hydrophobic groove formed by BH3, BH1, and BH2 domains.^21^ In addition, residues from each pocket are adjacent to or directly included in the Venetoclax drug’s binding site through the hydrophobic α4, α5, and α6 helices.^23^

## 4. Discussion

The significant differences in structural and functional metrics between HA sites and residue-matched random sites indicates HA site selection is not driven by residue type. Rather, we find common HA site amino acid types correspond with their structural and functional role. We note a high frequency of hydrophobic residues lysine, phenylalanine, and valine, corresponding with the “buried” position of HA sites near protein centroids. We also find a high frequency of glycine and cysteine, amino acids frequently found in tight turns and disulfide bonds respectively, indicating the critical functionality of HA sites. The low frequency of acidic and/or polar amino acids, including glutamic acid, tyrosine, and glutamine also matches our data, suggesting HA sites are rarely found on the protein surface and rather occupy deeper pockets.

While pairwise significant differences across the structural and functional metrics of primary, secondary, and tertiary HA sites are observed, these differences are less visually distinct than their respective comparisons to randomly selected sites, and we suspect this result is largely driven by high sample size and attention to outliers. We do observe a significant difference in the proportion of each HA site category across primary, secondary, and tertiary sites, though a weak association between HA site type and cluster assignment (p < 0.001, Cramer’s V = 0.051 with Chi-squared analysis, **Supplementary fig. 10**). Since primary HA sites are discovered first and have the highest proportion of core pathogenic sites and lowest proportion of low confidence sites, this suggests ESM-2 pays more attention to these functionally critical sites (i.e., core pathogenic sites) than sites it does not know how to characterize (i.e., low-confidence sites); however, the low count of low confidence sites across the groups limits this interpretation.

We conclude that using clustering-based categories is a more useful designation between HA sites. For low-confidence HA sites, we suspect ESM-2 pays high attention to these sites as it tries to understand their complex orientation relative to the other residues; with future generations of PLMs these sites may be better characterized, but in our method, they are indicated for filtering. Structural core HA sites, with proximity to protein centroid and known functional sites but low pathogenicity, appear important for defining the protein’s structural scaffolding and stable functions; that is, even when mutated, these HA sites will not significantly disturb the function of the functional sites they are close to. By contrast, structural pathogenic sites have highly pathogenic implications when mutated despite being distant from the protein centroid and known functional sites; mutations at these positions would significantly disrupt the structural stability, leading to pathogenic functional deficits elsewhere in the protein. Finally, the core pathogenic HA sites are close to the protein centroid and known functional sites *and* are strongly pathogenic when mutated. We thus interpret these sites as essential for regular protein structure and function.

One valuable use case of HA sites is for predicting pathogenic protein variants. Through improvements in genomic sequencing, identifying protein variants is a streamlined task, but interpreting their effect is less straightforward.^24^ Here, we show HA sites can be used to predict mutation pathogenicity. Our results using AlphaMissense predictions and real-world validation with PCAWG align with our understanding that HA sites have critical structural and functional roles, which when disrupted, may lead to disease. Further filtering to specifically explore structural pathogenic and core pathogenic HA sites could be used to predict downstream pathogenicity of previously undocumented variants, either in a clinical prognosis context or to anticipate the effects of designed protein alterations.

A second important use of HA sites is to identify candidate binding sites on proteins. In this study, we characterized the HA sites of 19,871 proteins; of these, less than a quarter (n=4,451 proteins; 22.4%) had UniProt-documented binding sites.^11^ The experimental process for identifying new binding sites with therapeutic benefit is a two-sided problem.^18^ First, a protein must be sufficiently characterized to identify candidate binding sites. Then, these sites must be screened with many drugs. Computational methods to narrow the field of candidate binding sites reduce the dimensionality of this procedure. Based on our results suggesting potential binding sites on two classically “undruggable” proteins, the transcription factor MYC and the anti-apoptotic protein BCL-2^18^, we propose HA sites could be used to narrow the field of candidate binding sites and thus improve the efficiency of both in vivo and in silico experiments.

We note the following limitations. We focus only on the ESM model, and while we expect other models with similar bidirectional structure to function similarly, their parameters may result in slight variations. We also use only one version of the ESM model, while other versions with different parameter specifications could influence high attention site specificity and categorization. Furthermore, we only calculate up to the third level of HA site, but additional levels of HA sites could be explored. We predict these would similarly categorize with clustering. However, we also note there is likely a natural limit to the level of HA site that can be determined, based on visual inspection of the attention matrices and considering the buffer used to generate sufficiently distinct HA site locations in each class.

## 5. Appendix

The code, data, and supplementary material accompanying this paper is available at https://github.com/Helix-Research-Lab/HAsite_fn.

