## Supplementary Figures for "Interpreting Protein Language Models: high attention sites predict functional regions"

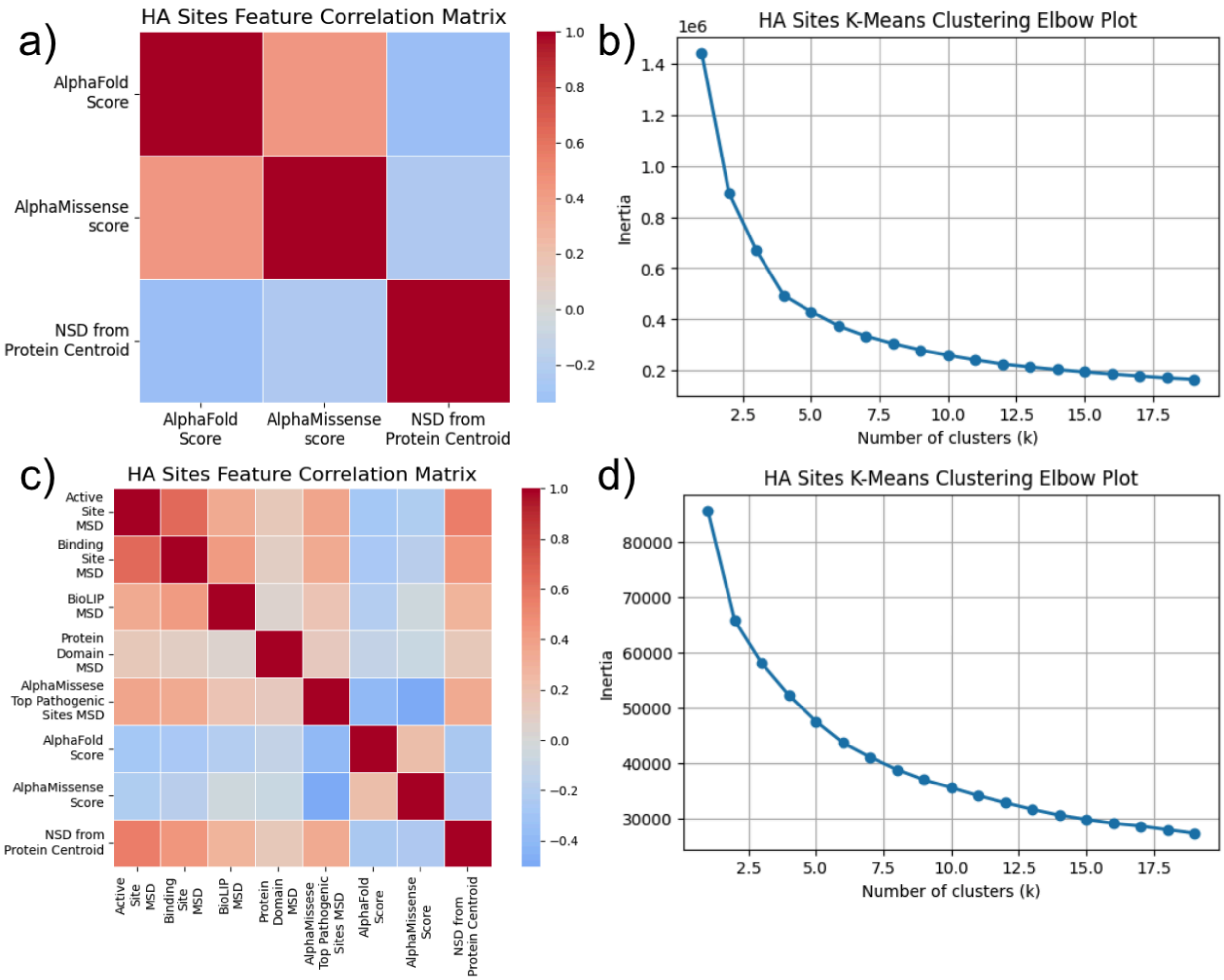


**Supplementary figure 1: HA sites clustering feature correlation analysis and parameter selection.** a) The Pearson correlation between reduced metric subset used to cluster HA sites. b) Elbow plot shows clustering stability (inertia) against number of clusters for K-means clustering of HA sites based on features in a). c) The Pearson correlation between extended metric set used to cluster HA sites. d) Elbow plot shows clustering stability (inertia) against number of clusters for K-means clustering of HA sites based on features in c).

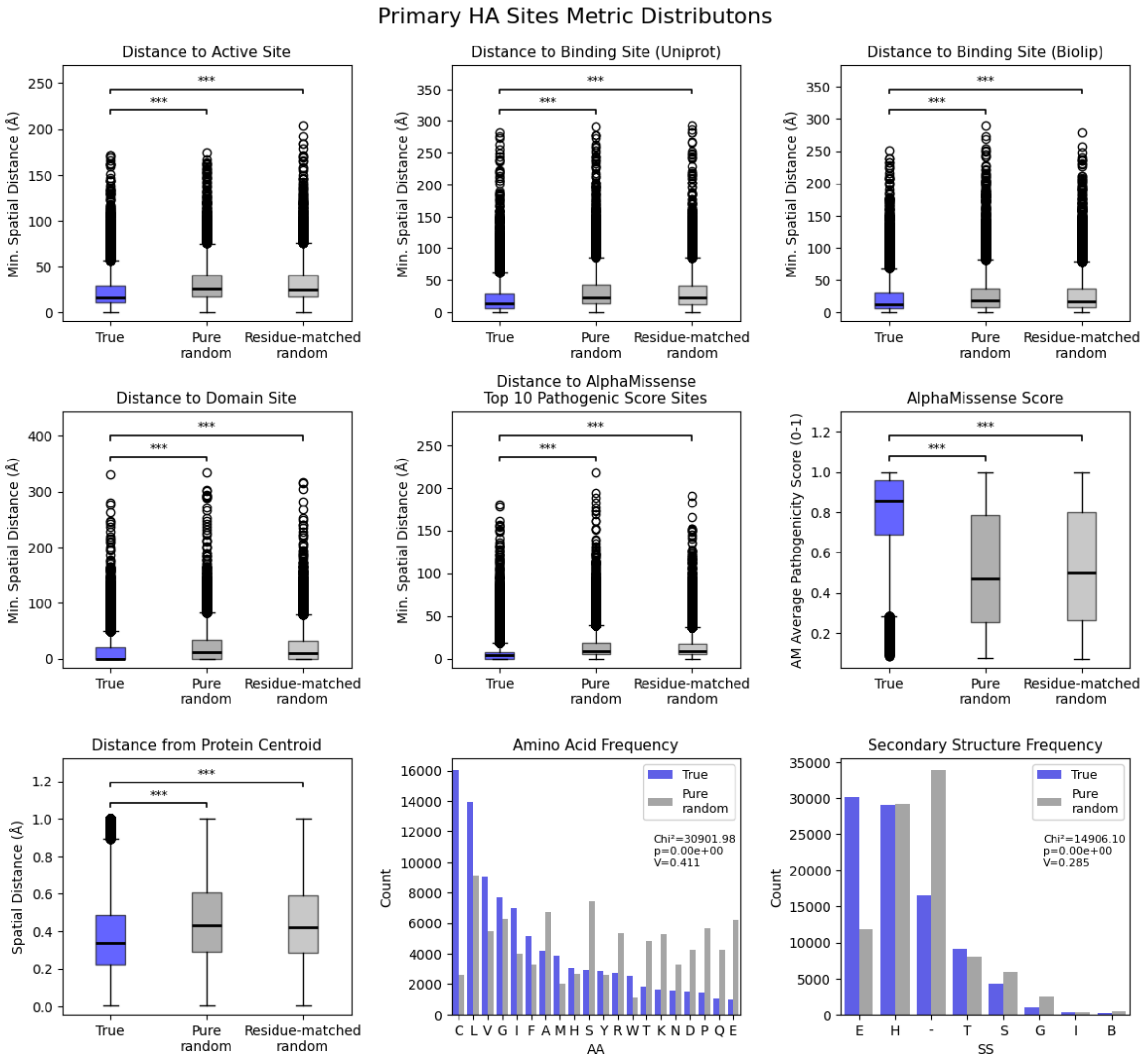


**Supplementary figure 2: Primary high attention site structural and functional metric distributions.** To evaluate the biological relevance of the primary HA sites, we plot structural and functional metrics of the primary HA sites (methods 2.3). Structural metrics include spatial orientation of the HA sites, described by the minimum spatial distance from HA sites to known active sites, binding sites, and domain sites (from UniProt and Biolip documentations), as well as the calculated protein centroid. Structural metrics also include HA site primary and secondary structure, shown as frequency plots of the amino acid type of HA sites and the secondary structures HA sites are part of. Functional metrics focus on pathogenicity of HA sites as predicted by AlphaMissense; the minimum distance from HA sites to the top 10 pathogenic sites in the protein as predicted by AlphaMissense, as well as the AlphaMissense score at the HA site, are shown. In blue we plot the metrics calculated at true HA sites. In dark grey, we plot the metrics calculated at randomized positions in the protein. In light grey, we plot the metrics calculated at residue-matched random positions (i.e., positions randomly selected from all residues of the same amino acid type as a given HA site). Two-sided Mann-Whitney tests with Bonferroni correction and Chi-squared analysis used to assess statistical significance. ***

indicates p-value < 0.001. V = Cramer's V. Amino acid single letter codes match standard definitions. Secondary structure single letter codes are defined as follows: E = extended strand, participates in β ladder; H = α-helix; - = coil / irregular (no defined secondary structure); T = hydrogen-bonded turn, S = bend, P = κ-helix (poly-proline II helix), G = 310-helix; I = π-helix; B = residue in isolated β-bridge. Across structural and functional metrics, we highlight the proximity of true primary HA sites to active and binding sites, as well as their significantly enriched pathogenicity (as calculated by the AlphaMissense score).

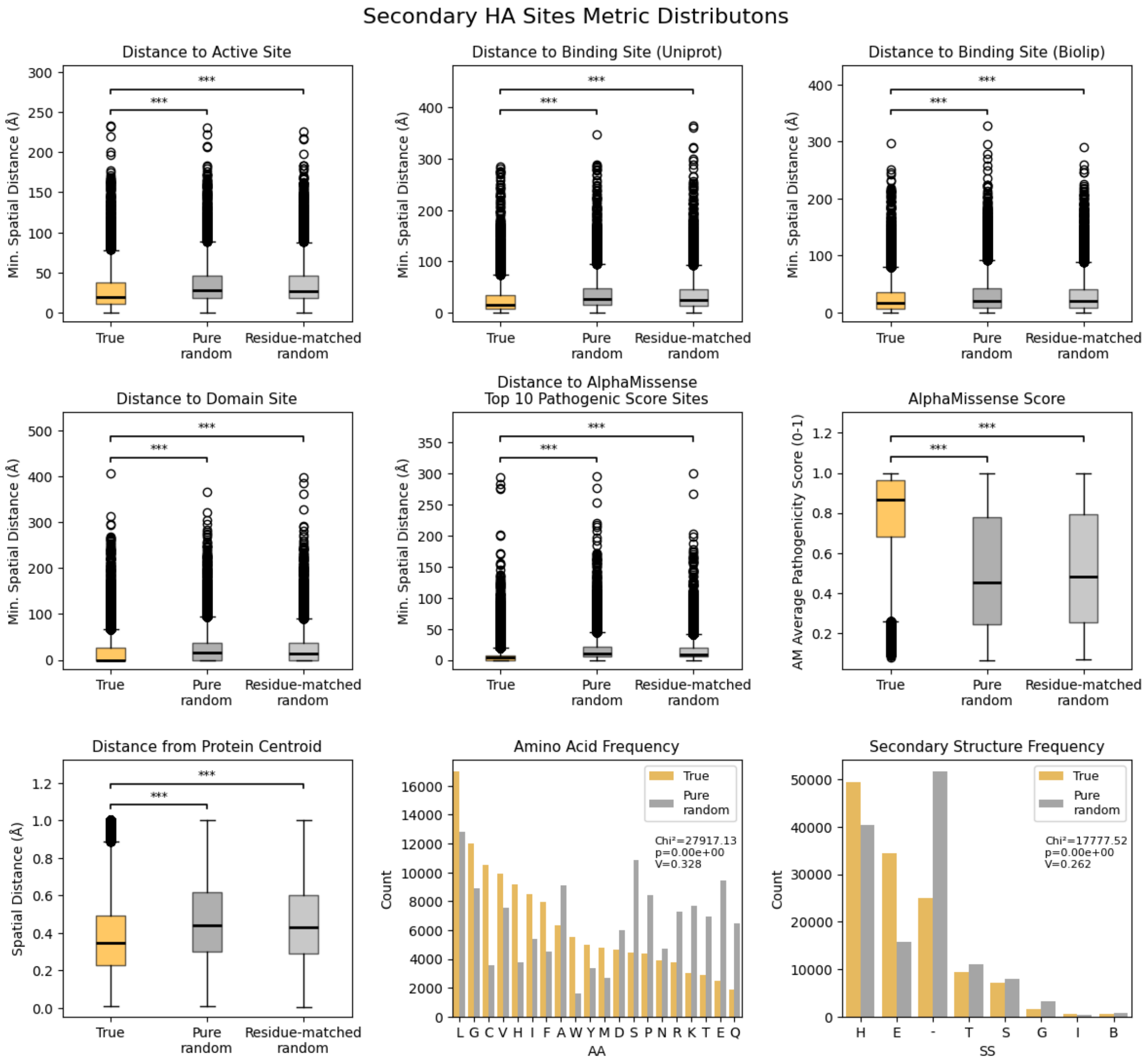


**Supplementary figure 3: Secondary high attention site structural and functional metric distributions.** To evaluate the biological relevance of the secondary HA sites, we plot structural and functional metrics of the secondary HA sites (methods 2.3). Structural metrics include spatial orientation of the HA sites, described by the minimum spatial distance from HA sites to known active sites, binding sites, and domain sites (from UniProt and Biolip documentations), as well as the calculated protein centroid. Structural metrics also include HA site primary and secondary structure, shown as frequency plots of the amino acid type of HA sites and the secondary structures HA sites are part of. Functional metrics focus on pathogenicity of HA sites as predicted by AlphaMissense; the minimum distance from HA sites to the top 10 pathogenic sites in the protein as predicted by AlphaMissense, as well as the AlphaMissense score at the HA site, are shown. In blue we plot the metrics calculated at true HA sites. In dark grey, we plot the metrics calculated at randomized positions in the protein. In light grey, we plot the metrics calculated at residue-matched random positions (i.e., positions randomly selected from all residues of the same amino acid type as a given HA site). Two-sided Mann-Whitney tests with

Bonferroni correction and Chi-squared analysis used to assess statistical significance. *** indicates p-value < 0.001. V = Cramer's V. Amino acid single letter codes match standard definitions. Secondary structure single letter codes are defined as follows: E = extended strand, participates in β ladder; H = α-helix; - = coil / irregular (no defined secondary structure); T = hydrogen-bonded turn, S = bend, P = κ-helix (poly-proline II helix), G = 310-helix; I = π-helix; B = residue in isolated β-bridge. Across structural and functional metrics, we highlight the proximity of true secondary HA sites to active and binding sites, as well as their significantly enriched pathogenicity (as calculated by the AlphaMissense score).

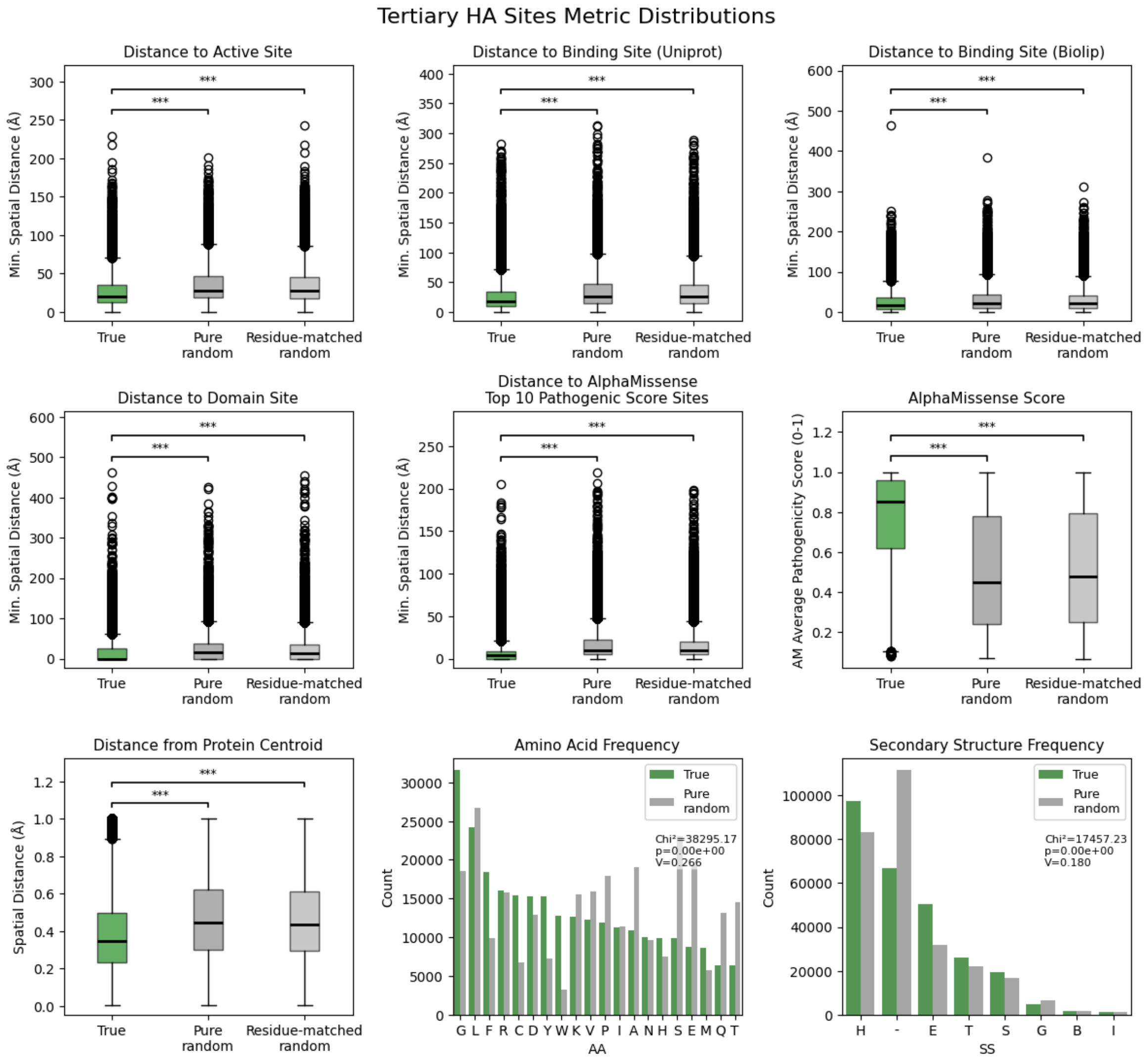


**Supplementary figure 4: Tertiary high attention site structural and functional metric distributions.** To evaluate the biological relevance of the tertiary HA sites, we plot structural and functional metrics of the tertiary HA sites (methods 2.3). Structural metrics include spatial orientation of the HA sites, described by the minimum spatial distance from HA sites to known active sites, binding sites, and domain sites (from UniProt and Biolip documentations), as well as the calculated protein centroid. Structural metrics also include HA site primary and secondary structure, shown as frequency plots of the amino acid type of HA sites and the secondary structures HA sites are part of. Functional metrics focus on pathogenicity of HA sites as predicted by AlphaMissense; the minimum distance from HA sites to the top 10 pathogenic sites in the protein as predicted by AlphaMissense, as well as the AlphaMissense score at the HA site, are shown. In blue we plot the metrics calculated at true HA sites. In dark grey, we plot the metrics calculated at randomized positions in the protein. In light grey, we plot the metrics calculated at residue-matched random positions (i.e., positions randomly selected from all residues of the same amino acid type as a given HA site). Two-sided Mann-Whitney tests with Bonferroni correction and Chi-squared analysis used to assess statistical significance. ***

indicates p-value < 0.001. V = Cramer's V. Amino acid single letter codes match standard definitions. Secondary structure single letter codes are defined as follows: E = extended strand, participates in β ladder; H = α-helix; - = coil / irregular (no defined secondary structure); T = hydrogen-bonded turn, S = bend, P = κ-helix (poly-proline II helix), G = 310-helix; I = π-helix; B = residue in isolated β-bridge. Across structural and functional metrics, we highlight the proximity of true tertiary HA sites to active and binding sites, as well as their significantly enriched pathogenicity (as calculated by the AlphaMissense score).

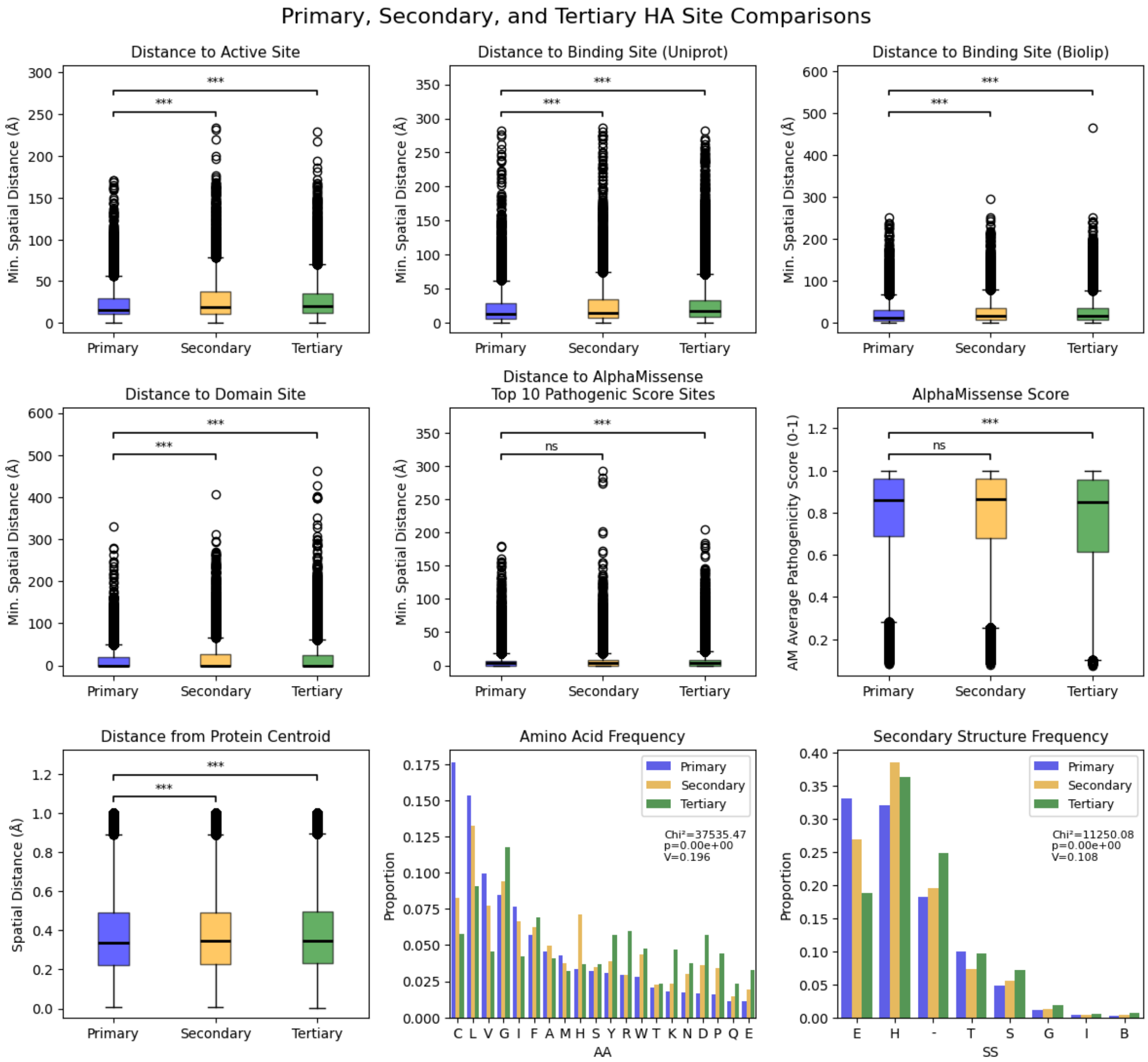


**Supplementary figure 5: Primary, secondary, and tertiary high attention site structural and functional metric distributions.** To evaluate the distinction between structural and functional metrics across primary, secondary, and tertiary HA sites, we plot structural and functional metrics of the primary, secondary, and tertiary HA sites against each other (methods 2.3). Structural metrics include spatial orientation of the HA sites, described by the minimum spatial distance from HA sites to known active sites, binding sites, and domain sites (from UniProt and Biolip documentations), as well as the calculated protein centroid. Structural metrics also include HA site primary and secondary structure, shown as frequency plots of the amino acid type of HA sites and the secondary structures HA sites are part of. Functional metrics focus on pathogenicity of HA sites as predicted by AlphaMissense; the minimum distance from HA sites to the top 10 pathogenic sites in the protein as predicted by AlphaMissense, as well as the AlphaMissense score at the HA site, are shown. In blue we plot the metrics calculated at primary HA sites; in orange, metrics calculated at secondary HA sites; in green, metrics calculated at tertiary HA sites. Two-sided Mann-Whitney tests with Bonferroni correction and Chi-squared analysis used to assess statistical significance. *** indicates p-value < 0.001. V = Cramer's V. Amino acid single letter codes match standard definitions. Secondary structure single letter codes are defined as follows: E = extended strand, participates in β ladder; H = α-helix; - = coil / irregular (no defined secondary structure); T = hydrogen-bonded turn, S = bend, P = κ-helix (poly-proline II helix), G = 310-helix; I = π-helix; B = residue in isolated β-bridge. Across structural and functional metrics, we highlight the proximity of true HA sites to active and binding sites, as well as their significantly enriched pathogenicity (as calculated by the AlphaMissense score). We highlight the relative similarity between primary, secondary, and tertiary HA sites across the metrics, though primary HA sites show greater proximity to active sites, binding sites, protein domains, and protein centroids, and tertiary sites show reduced pathogenicity as defined by AlphaMissense score.

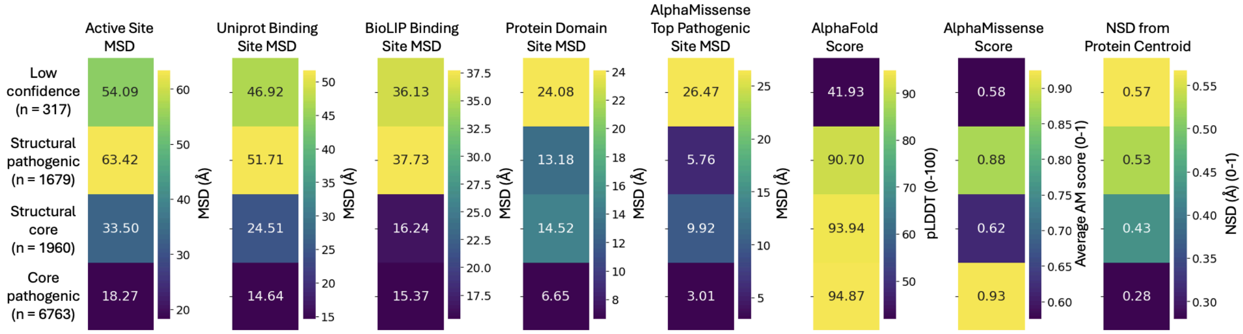


**Supplementary figure 8: Unsupervised clustering on extended features reveals four categories of HA sites.** Clustering performed on extended metric set (active site MSD, UniProt binding site MSD, BioLIP binding site MSD, protein domain site MSD, AlphaMissense top pathogenic site MSD, AlphaFold Score, AlphaMissense Score, NSD from Protein centroid) calculated per HA site (pooled primary, secondary, and tertiary). K-means clustering was used, returning an optimal K = 4 (methods 3.2). Active Site MSD = minimum spatial distance to known active sites; Uniprot Binding Site MSD = minimum spatial distance to known binding sites from Uniprot database; BioLIP Binding Site MSD = minimum spatial distance to known bidning sites from BioLIP database; Protein Domain Site MSD = minimum spatial distance to known domain sites from UniProt database; AlphaMissense Top Pathogenic Site MSD = minimum spatial distance to top 10 pathogenic sites per protein as defined by AlphaMissense score; AlphaFold Score = predicted local distance difference test (pLDDT) score; AlphaMissense Score = pathogenicity score; NSD from Protein Centroid = normalized spatial distance in Angstroms from protein centroid. Values in box represent average metric value for cluster. Left column counts represent number of HA sites in each cluster. Only HA sites from proteins with known active sites, binding sites, and protein domains are included in this clustering. Clustering with extended metric set produces the same four categories of HA sites, but shows additional metric variations explored in figure 5.

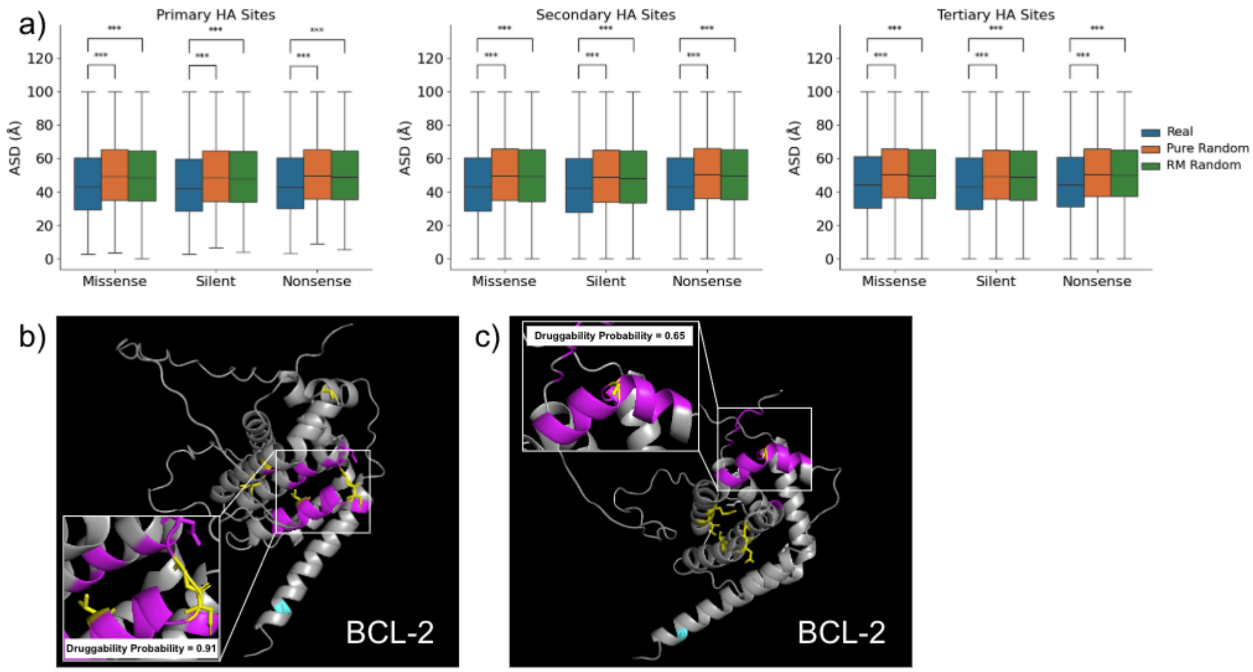


**Supplementary figure 9: HA sites are enriched near pathogenic variants and define novel binding sites.** a) Boxplot shows the average spatial distance (ASD) in Angstroms between each HA site residue and the nearest pathogenic variant (missense, nonsense, or silent) documented in the PCAWG cohort, separated by primary, secondary, and tertiary HA sites. Calculated distribution is plotted in blue and compared to both sets of randomized sites (methods). Two-sided Mann-Whitney tests with Bonferroni correction used to assess statistical significance. *** indicates p-value < 0.001. b) Pymol visualization of the BCL-2 protein (AlphaFold-generated structure) with HA sites highlighted. Yellow = core pathogenic HA sites, cyan = low-confidence HA sites. Pink = pocket of 5Å around core pathogenic HA sites. fPocket-predicted pocket druggability probability is shown. c) Pymol visualization of the BCL-2 protein (AlphaFold-generated structure) with HA sites highlighted. Yellow = core pathogenic HA sites, cyan = low-confidence HA sites. Pink = pocket of 8Å around core pathogenic HA site. fPocket-predicted pocket druggability probability is shown.

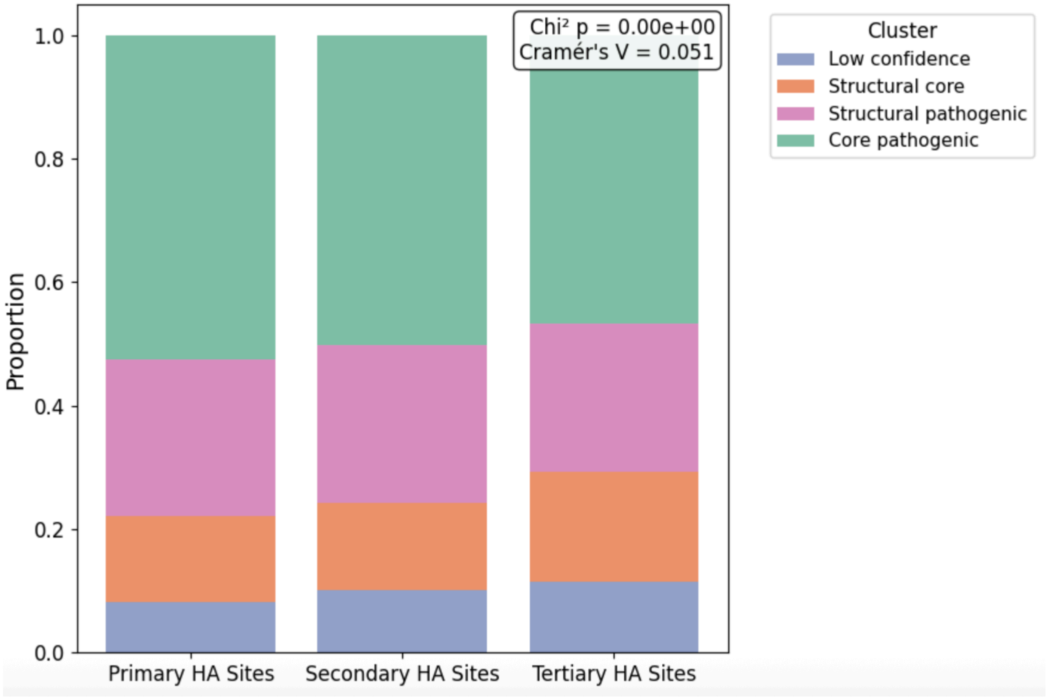


**Supplementary figure 10: Proportion of HA site type across primary, secondary, and tertiary HA sites.** Cluster assignments taken from main text (condensed) clustering on AlphaFold score, AlphaMissense score, and Protein Centroid NSD. Significance assessed with Chi-squared test. Primary, secondary, and tertiary HA sites are not strongly associated with HA site cluster type; all four categories of HA sites are seen across primary, secondary, and tertiary HA sites.
